# Identifying risk factors involved in the common versus specific liabilities to substance abuse: A genetically informed approach

**DOI:** 10.1101/595728

**Authors:** Eleonora Iob, Tabea Schoeler, Charlotte M. Cecil, Esther Walton, Andrew McQuillin, Jean-Baptiste Pingault

## Abstract

The co-occurrence of abuse of multiple substances is thought to stem from a common liability that is partly genetic in origin. Genetic risk may indirectly contribute to a common liability through genetically influenced individual vulnerabilities and traits. To disentangle the aetiology of common versus specific liabilities to substance abuse, polygenic scores can be used as genetic proxies indexing such risk and protective individual vulnerabilities or traits. In this study, we used genomic data from a UK birth cohort study (ALSPAC, N*=*4218) to generate 18 polygenic scores indexing mental health vulnerabilities, personality traits, cognition, physical traits, and substance abuse. Common and substance-specific factors were identified based on four classes of substance abuse (alcohol, cigarettes, cannabis, other illicit substances) assessed over time (age 17, 20, and 22). In multivariable regressions, we then tested the independent contribution of selected polygenic scores to the common and substance-specific factors. Our findings implicated several genetically influenced traits and vulnerabilities in the common liability to substance abuse, most notably risk taking (*b*_standardized_=0.14; 95%CI: 0.10,0.17), followed by extraversion (*b*_standardized_ =-0.10; 95%CI: −0.13,-0.06), and schizophrenia risk (*b*_standardized_=0.06; 95%CI: 0.02;0.09). Educational attainment (EA) and body mass index (BMI) had opposite effects on substance-specific liabilities such as cigarettes (*b*_standardized-EA_= −0.15; 95%CI: −0.19,-0.12; *b*_standardized-BMI_=0.05; 95%CI: 0.02,0.09), alcohol (*b*_standardized-EA_=0.07; 95%CI: 0.03,0.11; *b*_standardized-BMI_= −0.06; 95%CI: −0.10, −0.02), and other illicit substances (*b*_standardized-EA_=0.12; 95%CI: 0.07,0.17; *b*_standardized-BMI_= −0.08; 95%CI:-0.13,-0.04). This is the first study based on genomic data that clarifies the aetiological architecture underlying the common versus substance-specific liabilities, providing novel insights for the prevention and treatment of substance abuse.

## INTRODUCTION

Substance abuse is a leading contributor to the global disease and disability burden^1^ and associated with high societal and economic costs. Observational studies consistently report substantial correlations between the abuse of distinct substances such as cigarettes, alcohol and cannabis^2-6^. High rates of abuse of multiple substances are of particular public health concern given the pervasive long-term consequences on health of such pattern of co-occurrence^6-8^. According to the common liability model, the observed correlations between the abuse of distinct substances can be explained by the presence of a common, nonspecific liability underlying different classes of substances^9^. Support for this model comes from several lines of research. For example, in observational studies, the use of different classes of substances is typically associated with a range of shared individual factors such as mental health vulnerabilities [e.g. schizophrenia, attention deficit and hyperactivity disorder (ADHD)]^10-12^, personality traits (e.g. risk-taking)^13-15^, and cognitive factors (e.g. low educational attainment)^16,17^. Results from twin^3,18^ and genomic studies^19,20^ further indicate that the correlation between the use of different substances stems from a common liability that is largely genetic in nature.

A theory of a common (genetic) liability is, however, somewhat conflicting with findings from genome-wide association studies (GWAS). Up to now, single nucleotide polymorphisms (SNPs) identified in GWAS are mainly associated with the use of particular classes of substances^20-23^. For example, a replicated finding is the association between the alcohol metabolism gene Alcohol Dehydrogenase 1B (ADH1B) and alcohol abuse^20^, or the association between the nicotine metabolism gene Cholinergic Receptor Nicotinic Alpha 5 Subunit (CHRNA5) and nicotine use^20^. One reason why studies have yet failed to uncover genetic influences on the common liability may lie in the fact that, so far, GWAS have not systematically modelled factors that reflect common and substance-specific liabilities. Moreover, the genetic architecture of the common liability may consist of highly polygenic and small indirect effects via a range of genetically influenced traits and vulnerabilities, such as depression or risk taking. For example, many genetic variants influence risk taking, which in turn may contribute to the common liability. As such, if those traits and vulnerabilities are causally involved in the aetiology of the common liability to substance abuse, their respective genetic proxies (e.g. genetic variants associated with risk taking) must be associated with the common liability.

In this study, we propose to exploit the polygenic score (PGS) approach to further interrogate the aetiology of the common and substance-specific liabilities to substance abuse. A polygenic score is a continuous index of an individual’s genetic risk for a particular phenotype, based on GWAS results for the corresponding phenotype^24^. PGSs can be used as genetic proxies indexing vulnerabilities and traits to study their role in the common and specific liabilities to substance abuse. Employing PGSs as proxies for potential risk factors can be conceived as a first step in a series of genetically-informed designs to strengthen causal evidence in observational studies^25,26^. For example, studies have used PGSs indexing a particular vulnerability or trait, such as depression or psychotic disorders, to test their association with the use of specific classes of substances including cannabis^27^, alcohol^28,29^, nicotine^28,29^ or illicit substances^28^. However, this evidence does not provide insights regarding the aetiology of common versus substance-specific liabilities. One study has employed the PGS approach to study the effect of a few selected PGSs indexing mental health disorders on the abuse of multiple substances^30^. However, important traits and vulnerabilities previously implicated in the aetiology of substance abuse, including personality traits and cognitive measures, remain as yet unexplored.

In this study, we aimed to triangulate and extend previous phenotypic evidence on risk and protective factors for the common versus specific liabilities to substance abuse by integrating genomic data from a longitudinal population-based cohort. Specifically, we generated 18 PGSs, reflecting a range of individual mental health vulnerabilities and traits previously implicated in the aetiology of substance use. Using the PGS approach, we tested their associations with (1) a common liability to substance abuse capturing the co-occurrence of abuse of alcohol, cigarettes, cannabis, and other illicit substances, as well as (2) substance-specific liabilities that are independent of the common liability. By applying genetically informed methods to study refined phenotypes, this investigation has the potential to yield important insights for the aetiology of substance abuse and inform prevention and treatment programs.

## METHODS

### Sample

We analysed data from the Avon Longitudinal Study of Parents and Children (ALSPAC)^31^. Details about the study design, methods of data collection, and variables can be found on the study website (http://www.bristol.ac.uk/alspac/researchers/our-data/). We used phenotypic data on substance abuse collected when the study participants were aged 17, 20, and 22 years. Genotype data was available for 7288 unrelated children of European ancestry after quality control [cf. Supplementary Information (SI) for details]. After excluding individuals without sufficient data on substance abuse, the final sample consisted of 4218 individuals. Included individuals differed from non-individuals in several sample characteristics (eTable1, SI), but differences were only small in magnitude. Ethical approval for the study was obtained from the ALSPAC Ethics and Law Committee and the Local Research Ethics Committees.

**Table 1.** Descriptive statistics of the four substance abuse measures at age 17, 20, and 22.

|  |  | Age 17 (%) | Age 20 (%) | Age 22 (%) |
| --- | --- | --- | --- | --- |
| **Alcohol**<br>(Audit) | *Non-users* | 4.7 | 2.0 | 5.0 |
|  | *Low Abuse* | 54.5 | 41.8 | 52.0 |
|  | *Moderate Abuse* | 36.7 | 42.9 | 37.2 |
|  | *High Abuse* | 4.1 | 13.3 | 5.8 |
|  | *Missing* | 32.7 | 36.3 | 38.1 |
| **Cigarettes**<br>(Fagerstrom) | *Non-users* | 48.5 | 39.6 | 39.5 |
|  | *Occasional users* | 39.6 | 42.5 | 48.6 |
|  | *Low Abuse* | 4.8 | 10.0 | 5.6 |
|  | *Moderate Abuse* | 4.5 | 5.4 | 4.5 |
|  | *High Abuse* | 2.5 | 2.6 | 1.7 |
|  | *Missing* | 30.9 | 32.2 | 37.3 |
| **Cannabis**<br>(Cast) | *Non-users* | 92.4 | 84.6 | 88.6 |
|  | *Low Abuse* | 5.2 | 12.6 | 9.4 |
|  | *Moderate Abuse* | 2.1 | 2.4 | 1.9 |
|  | *High Abuse* | 0.4 | 0.4 | 0.2 |
|  | *Missing* | 39.7 | 32.4 | 37.7 |
| **Other substance** | *Non-users* | 85.8 | 80.0 | 72.6 |
|  | *Low Abuse* | 10.9 | 14.5 | 18.5 |
|  | *Moderate Abuse* | 2.6 | 4.4 | 7.0 |
|  | *High Abuse* | 0.7 | 1.2 | 1.9 |
|  | *Missing* | 32.1 | 34.9 | 39.9 |

**Note.** Data source = ALSPAC. The total number of non-users and users is 100%. Missing values refer to the initial sample of genotyped participants (N = 7288) leading to a total of 4218 participants with at least one substance use measure across all time points, who were included in the structural equation model. Audit = Alcohol Use Disorders Identification Test. Cast = Cannabis Abuse Screening Test. Fagerstrom = Fagerstrom Test of Nicotine Dependence. Other substance = total number of other illicit substances consumed in the past 12 months (i.e. cocaine, ketamine, heroin etc.) (cf. SI).

### Measures

#### Substance abuse

Substance abuse (i.e. cigarette, alcohol, cannabis, and other illicit substances) was measured at ages 17, 20 and 22 using validated self-report questionnaires. These included: the Fagerstrom Test for Nicotine Dependence^32^, the Alcohol Use Disorders Identification Test^33^, and the Cannabis Abuse Screening Test^34^. For each scale, total scores were calculated by adding up their item scores. For other illicit substance abuse, we computed the total number of illicit substances used in the previous 12 months (cf. SI for details).

#### Summary Statistics Datasets

We collected summary statistics from 34 publicly available GWAS derived from discovery cohorts which did not include ALSPAC participants (eTable 2, SI), indexing domains such as mental health vulnerabilities (e.g. depression), personality (e.g. risk taking), cognition (e.g. educational attainment), physical measures [e.g. body mass index (BMI)], and substance use (i.e. nicotine, alcohol, and cannabis use). From the initial 34 GWAS, we excluded several GWAS to avoid multicollinearity and increase power, resulting in a final selection of 18 GWAS summary statistics (details on exclusion criteria in SI, eTable 3).

**Table 2.** Single-PGS TSO models.

| PGSs | Common Substance Abuse |  |  |  |  | Cigarette Abuse |  |  |  |  | Alcohol Abuse |  |  |  |  | Cannabis Abuse |  |  |  |  | Other Substance Abuse |  |  |  |  |
| --- | --- | --- | --- | --- | --- | --- | --- | --- | --- | --- | --- | --- | --- | --- | --- | --- | --- | --- | --- | --- | --- | --- | --- | --- | --- |
|  | Coef. | 95% CI | Stand. coef. | p | p (perm/FDR) | Coef. | 95% CI | Stand. Coef. | p | p (perm/FDR) | Coef. | 95% CI | Stand. Coef. | p | p (perm/FDR) | Coef. | 95% CI | Stand. Coef. | p | p (perm/FDR) | Coef. | 95% CI | Stand. Coef. | p | p (perm/FDR) |
| Substance Abuse |  |  |  |  |  |  |  |  |  |  |  |  |  |  |  |  |  |  |  |  |  |  |  |  |  |
| Cigarettes Per Day | 0.015 | -0.007; 0.037 | 0.028 | 0.193 | 0.313 | 0.122 | 0.094; 0.15 | 0.168 | <.001 | <.001 | -0.023 | -0.048; 0.003 | -0.035 | 0.087 | 0.182 | 0.009 | -0.017; 0.035 | 0.017 | 0.496 | 0.603 | -0.050 | -0.072; -0.028 | -0.11 | <.001 | <.001 |
| Cigarettes Age Onset | -0.066 | -0.087; -0.044 | -0.124 | <.001 | <.001 | -0.098 | -0.129; -0.067 | -0.134 | <.001 | <.001 | 0.018 | -0.007; 0.044 | 0.029 | 0.155 | 0.263 | -0.049 | -0.074; -0.024 | -0.094 | <.001 | <.001 | 0.038 | 0.015; 0.062 | 0.085 | 0.001 | 0.004 |
| Alcohol Dependence | 0.025 | 0.005; 0.045 | 0.047 | 0.015 | 0.045 | 0.028 | 0.002; 0.053 | 0.038 | 0.034 | 0.085 | 0.029 | 0.005; 0.053 | 0.045 | 0.019 | 0.052 | 0.010 | -0.012; 0.032 | 0.019 | 0.382 | 0.491 | -0.013 | -0.034; 0.007 | -0.029 | 0.198 | 0.313 |
| Alcohol Per Week | 0.099 | 0.078; 0.119 | 0.186 | <.001 | <.001 | -0.015 | -0.039; 0.009 | -0.020 | 0.231 | 0.352 | 0.219 | 0.197; 0.242 | 0.343 | <.001 | <.001 | -0.033 | -0.055; -0.012 | -0.066 | 0.002 | 0.007 | 0.051 | 0.027; 0.074 | 0.108 | <.001 | <.001 |
| Cannabis Use Disorder | 0.011 | -0.009; 0.030 | 0.020 | 0.283 | 0.392 | -0.002 | -0.027; 0.023 | -0.003 | 0.872 | 0.922 | 0.038 | 0.013; 0.063 | 0.059 | 0.003 | 0.010 | 0.005 | -0.017; 0.027 | 0.009 | 0.681 | 0.796 | -0.019 | -0.041; 0.003 | -0.041 | 0.097 | 0.194 |
| Cannabis Use Frequency | 0.023 | 0.004; 0.043 | 0.044 | 0.019 | 0.052 | 0.001 | -0.023; 0.025 | 0.001 | 0.962 | 0.962 | 0.024 | 0.001; 0.048 | 0.038 | 0.041 | 0.100 | 0.008 | -0.013; 0.029 | 0.016 | 0.464 | 0.572 | -0.001 | -0.022; 0.019 | -0.003 | 0.909 | 0.922 |
| Mental Health |  |  |  |  |  |  |  |  |  |  |  |  |  |  |  |  |  |  |  |  |  |  |  |  |  |
| ADHD | 0.012 | -0.008; 0.032 | 0.022 | 0.242 | 0.360 | 0.047 | 0.022; 0.073 | 0.065 | <.001 | <.001 | -0.018 | -0.043; 0.006 | -0.029 | 0.141 | 0.254 | 0.024 | 0.002; 0.047 | 0.047 | 0.034 | 0.085 | -0.031 | -0.052; -0.009 | -0.067 | 0.005 | 0.016 |
| Depression | 0.021 | 0.000; 0.041 | 0.039 | 0.047 | 0.108 | 0.044 | 0.019; 0.070 | 0.061 | 0.001 | 0.004 | -0.010 | -0.035; 0.014 | -0.016 | 0.407 | 0.509 | 0.025 | 0.004; 0.047 | 0.049 | 0.019 | 0.052 | -0.031 | -0.053; -0.010 | -0.069 | 0.005 | 0.016 |
| Worry | -0.012 | -0.033; 0.009 | -0.023 | 0.250 | 0.360 | 0.006 | -0.019; 0.030 | 0.008 | 0.651 | 0.780 | -0.003 | -0.026; 0.021 | -0.005 | 0.810 | 0.900 | 0.003 | -0.019; 0.025 | 0.006 | 0.780 | 0.889 | -0.015 | -0.036; 0.006 | -0.033 | 0.154 | 0.263 |
| Schizophrenia | 0.038 | 0.019; 0.057 | 0.072 | <.001 | <.001 | -0.020 | -0.045; 0.004 | -0.028 | 0.099 | 0.194 | 0.041 | 0.016; 0.065 | 0.063 | 0.001 | 0.004 | 0.014 | -0.007; 0.035 | 0.027 | 0.196 | 0.313 | 0.018 | -0.003; 0.039 | 0.039 | 0.091 | 0.186 |
| Personality |  |  |  |  |  |  |  |  |  |  |  |  |  |  |  |  |  |  |  |  |  |  |  |  |  |
| Extraversion | -0.054 | -0.074; -0.034 | -0.102 | <.001 | <.001 | -0.017 | -0.043; 0.008 | -0.024 | 0.187 | 0.312 | -0.076 | -0.100; -0.051 | -0.117 | <.001 | <.001 | 0.022 | 0.001; 0.044 | 0.044 | 0.044 | 0.104 | -0.035 | -0.056; -0.014 | -0.075 | 0.001 | 0.004 |
| Irritability | 0.012 | -0.007; 0.032 | 0.023 | 0.219 | 0.34 | 0.020 | -0.005; 0.046 | 0.028 | 0.116 | 0.213 | 0.012 | -0.013; 0.036 | 0.018 | 0.348 | 0.461 | 0.018 | -0.004; 0.040 | 0.035 | 0.108 | 0.207 | -0.013 | -0.034; 0.009 | -0.028 | 0.252 | 0.360 |
| Neuroticism | -0.001 | -0.021; 0.019 | -0.002 | 0.903 | 0.922 | 0.011 | -0.014; 0.036 | 0.015 | 0.398 | 0.505 | 0.015 | -0.010; 0.040 | 0.023 | 0.252 | 0.360 | 0.002 | -0.019; 0.023 | 0.004 | 0.864 | 0.922 | -0.010 | -0.032; 0.012 | -0.022 | 0.361 | 0.471 |
| Risk Taking | 0.077 | 0.057; 0.097 | 0.145 | <.001 | <.001 | 0.033 | 0.009; 0.057 | 0.045 | 0.007 | 0.022 | 0.046 | 0.021; 0.070 | 0.071 | <.001 | <.001 | 0.017 | -0.006; 0.040 | 0.033 | 0.147 | 0.259 | 0.002 | -0.021; 0.025 | 0.004 | 0.884 | 0.922 |
| Physical Development |  |  |  |  |  |  |  |  |  |  |  |  |  |  |  |  |  |  |  |  |  |  |  |  |  |
| Birth weight | 0.001 | -0.018; 0.021 | 0.002 | 0.904 | 0.922 | -0.026 | -0.051; -0.002 | -0.036 | 0.034 | 0.085 | 0.003 | -0.022; 0.028 | 0.005 | 0.803 | 0.900 | 0.013 | -0.010; 0.036 | 0.026 | 0.264 | 0.371 | 0.005 | -0.018; 0.028 | 0.011 | 0.659 | 0.780 |
| BMI | -0.001 | -0.021; 0.019 | -0.002 | 0.912 | 0.922 | 0.067 | 0.042; 0.093 | 0.093 | <.001 | <.001 | -0.043 | -0.068; -0.018 | -0.067 | 0.001 | 0.004 | 0.021 | -0.001; 0.043 | 0.041 | 0.062 | 0.133 | -0.054 | -0.076; -0.032 | -0.117 | <.001 | <.001 |
| Height | 0.021 | 0.000; 0.041 | 0.039 | 0.049 | 0.110 | -0.013 | -0.037; 0.012 | -0.018 | 0.304 | 0.408 | 0.024 | -0.001; 0.049 | 0.038 | 0.057 | 0.125 | 0.017 | -0.004; 0.039 | 0.034 | 0.114 | 0.213 | 0.001 | -0.02; 0.023 | 0.003 | 0.904 | 0.922 |
| Cognition |  |  |  |  |  |  |  |  |  |  |  |  |  |  |  |  |  |  |  |  |  |  |  |  |  |
| Educational Attainment | 0.003 | -0.016; 0.022 | 0.006 | 0.736 | 0.849 | -0.122 | -0.149; -0.095 | -0.168 | <.001 | <.001 | 0.050 | 0.027; 0.074 | 0.078 | <.001 | <.001 | -0.012 | -0.033; 0.010 | -0.023 | 0.289 | 0.394 | 0.067 | 0.045; 0.09 | 0.146 | <.001 | <.001 |

**Note.** Data source: ALSPAC, N = 4218. Coef. = regression coefficient. Stand coef. = standardised regression coefficient. CI = confidence interval. p = p-value. p (perm/FDR) = False Discovery Rate (FDR) corrected p-values. Estimator = MLR (maximum likelihood estimation with robust standard errors). TSO = Trait-State-Occasion. PGS = Polygenic Score. ADHD = Attention deficit hyperactivity disorder. BMI = Body Mass Index.

**Table 3.** Multi-PGSs TSO models.

| Common Substance Abuse |  |  |  |  |  | Cigarette Abuse |  |  |  |  | Alcohol Abuse |  |  |  |  | Cannabis Abuse |  |  |  |  | Other Substance Abuse |  |  |  |  |
| --- | --- | --- | --- | --- | --- | --- | --- | --- | --- | --- | --- | --- | --- | --- | --- | --- | --- | --- | --- | --- | --- | --- | --- | --- | --- |
| PGSs | Coef. | 95% CI | Stand. Coef. | p | p (perm/FDR) | Coef. | 95% CI | Stand. Coef. | p | p (perm/FDR) | Coef. | 95% CI | Stand. Coef. | p | p (perm/FDR) | Coef. | 95% CI | Stand. Coef. | p | p (perm/FDR) | Coef. | 95% CI | Stand. Coef. | p | p (perm/FDR) |
| Substance Abuse |  |  |  |  |  |  |  |  |  |  |  |  |  |  |  |  |  |  |  |  |  |  |  |  |  |
| Cigarettes Per Day |  |  |  |  |  | 0.109 | 0.081; 0.137 | 0.150 | <.001 | <.001 |  |  |  |  |  |  |  |  |  |  | -0.048 | -0.071; -0.026 | -0.104 | <.001 | <.001 |
| Cigarettes Age Onset | -0.060 | -0.081; -0.039 | -0.108 | <.001 | <.001 | -0.079 | -0.109; -0.048 | -0.108 | <.001 | <.001 |  |  |  |  |  | -0.050 | -0.075; -0.025 | -0.096 | <.001 | <.001 | 0.034 | 0.010; 0.057 | 0.073 | 0.006 | 0.008 |
| Alcohol Dependence | 0.016 | -0.004; 0.036 | 0.028 | 0.122 | 0.131 |  |  |  |  |  |  |  |  |  |  |  |  |  |  |  |  |  |  |  |  |
| Alcohol Per Week | 0.094 | 0.073; 0.115 | 0.169 | <.001 | <.001 |  |  |  |  |  | 0.219 | 0.196; 0.241 | 0.342 | <.001 | <.001 | -0.035 | -0.056; -0.014 | -0.067 | 0.001 | 0.002 | 0.055 | 0.032; 0.078 | 0.119 | <.001 | <.001 |
| Cannabis Use Disorder |  |  |  |  |  |  |  |  |  |  | 0.030 | 0.006; 0.054 | 0.047 | 0.013 | 0.016 |  |  |  |  |  |  |  |  |  |  |
| Individual Vulnerability and Protective Factors |  |  |  |  |  |  |  |  |  |  |  |  |  |  |  |  |  |  |  |  |  |  |  |  |  |
| ADHD |  |  |  |  |  | 0.017 | -0.009; 0.043 | 0.023 | 0.210 | 0.217 |  |  |  |  |  |  |  |  |  |  | -0.014 | -0.036; 0.009 | -0.030 | 0.228 | 0.228 |
| Depression |  |  |  |  |  | 0.027 | 0.001; 0.052 | 0.037 | 0.043 | 0.050 |  |  |  |  |  |  |  |  |  |  | -0.020 | -0.043; 0.002 | -0.044 | 0.078 | 0.087 |
| Schizophrenia | 0.030 | 0.011; 0.049 | 0.056 | 0.002 | 0.003 |  |  |  |  |  | 0.032 | 0.007; 0.056 | 0.049 | 0.011 | 0.014 |  |  |  |  |  |  |  |  |  |  |
| Extraversion | -0.051 | -0.071; -0.030 | -0.095 | <.001 | <.001 |  |  |  |  |  | -0.076 | -0.100; -0.051 | -0.118 | <.001 | <.001 |  |  |  |  |  | -0.040 | -0.061; -0.018 | -0.085 | <.001 | <.001 |
| Risk Taking | 0.072 | 0.052; 0.092 | 0.136 | <.001 | <.001 | 0.035 | 0.011; 0.059 | 0.048 | 0.004 | 0.006 | 0.041 | 0.016; 0.065 | 0.063 | 0.001 | 0.002 |  |  |  |  |  |  |  |  |  |  |
| BMI |  |  |  |  |  | 0.038 | 0.011; 0.065 | 0.052 | 0.005 | 0.007 | -0.035 | -0.061; -0.010 | -0.055 | 0.007 | 0.009 |  |  |  |  |  | -0.039 | -0.062; -0.016 | -0.084 | 0.001 | 0.002 |
| Educational Attainment |  |  |  |  |  | -0.110 | -0.138; -0.083 | -0.151 | <.001 | <.001 | 0.044 | 0.019; 0.068 | 0.068 | <.001 | <.001 |  |  |  |  |  | 0.057 | 0.033; 0.080 | 0.121 | <.001 | <.001 |

**Note.** Data source: ALSPAC, N = 4218. Coef. = regression coefficient. Stand coef. = standardised regression coefficient. CI = confidence interval. p = p-value. p (perm/FDR) = False Discovery Rate (FDR) corrected p-values. Estimator = MLR (maximum likelihood estimation with robust standard errors). TSO = Trait-State-Occasion. PGS = Polygenic Score. ADHD = Attention deficit hyperactivity disorder. BMI = Body Mass Index.

### Statistical analyses

#### Polygenic score analyses

18 PGSs were generated utilising PRSice software (http://www.prsice.info/)^24^, based on ALSPAC genotype data and the selected GWAS summary statistics. The PGSs for each individual were calculated as the sum of alleles associated with the phenotype of interest (e.g. schizophrenia), weighted by their effect sizes found in the corresponding GWAS. Clumping was performed in order to remove SNPs in linkage disequilibrium (r^2^ > 0.10 within a 250-bp window). The PGSs were generated using a p-value threshold of 1.

#### Trait-State-Occasion models of substance abuse

All analyses were conducted in R version 3.5.1, using the ‘Lavaan’ package^35^. First, Trait-State-Occasion (TSO) structural equation models were fitted using the scores for cigarette, alcohol, cannabis, and other illicit substance abuse at each time point^36,37^. This approach enabled us to model latent factors of substance abuse that are stable over time, including (i) a common factor of all substances and (ii) substance-specific factors. Such advanced phenotypic modelling retains a higher degree of precision and specificity compared to simple observed substance use phenotypes. Missing data on the substance use indicators were handled using Full Maximum Likelihood Estimation. The model parameters were estimated using robust standard errors due to non-normality of the substance abuse scores. The TSO model was tested using available model specifications^38^. Further details are provided in the eMethods (SI) and in illustrative Figures [cf. Figure 1 (simplified illustration) and eFigure 1 (full specification, SI)]. Second, we tested the associations of each PGS with both the common and the substance-specific latent factors by incorporating regression analysis into the TSO models (single-PGS TSO models). False Discovery Rate (FDR) corrected p-values^39^ are provided to account for multiple testing. Finally, we tested two sets of multivariable TSO models (multi-PGSs TSO models) for each latent factor, in which we included only those PGSs that remained significant after FDR correction. In the first set, we included PGSs indexing substance use phenotypes (i.e. PGSs indexing dependency and frequency of cigarette, cannabis and alcohol use). In the second set, we included PGSs indexing mental health vulnerabilities and traits. All regression models were controlled for sex and population stratification by including 10 principal components as covariates. All PGSs were standardised.

**Figure 1.**
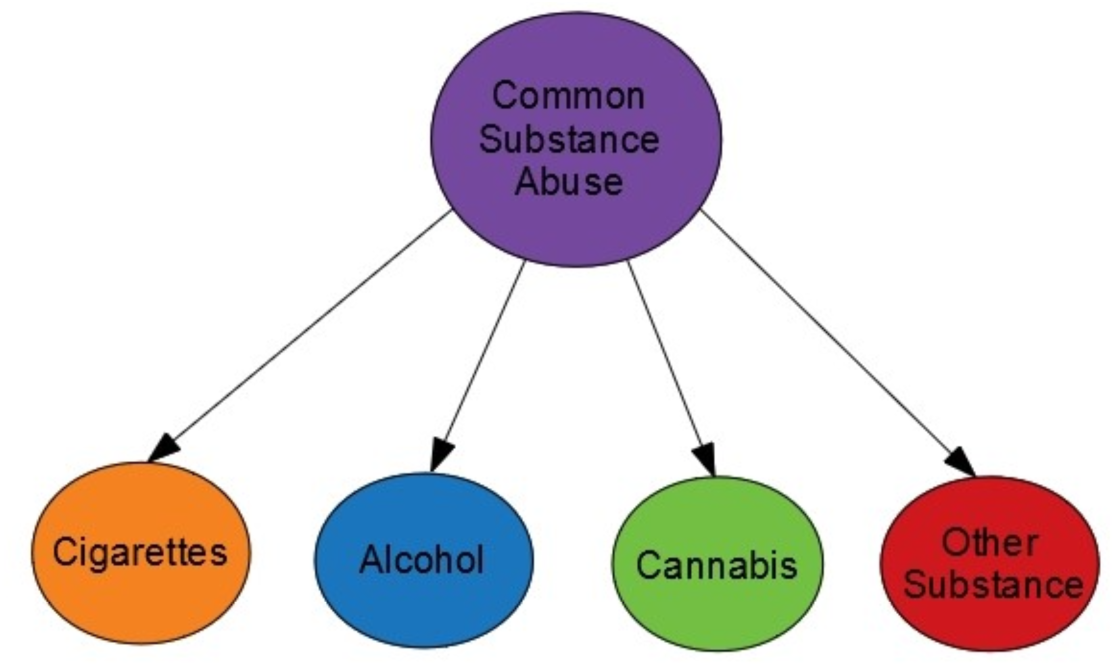
Modelling common versus specific liabilities to substance abuse. **Note.** Simplified illustration of the Trait-State-Occasion (TSO) model of substance abuse. The full TSO model includes modelling over three time points and is detailed in the SI (eFigure 1).

## RESULTS

The patterns of substance abuse in our sample are provided in Table 1. Correlations between the 18 PGSs and phenotypic measures of substance abuse are displayed in Figure 2 and provided in eTable 5 (SI). The Trait-State-Occasion (TSO) model of substance abuse fits the data reasonably well (*X*^*2*^ (42)=284.67, *p*<0.001, CFI=0.952, RMSEA=0.037, SRMR=0.058)^37^. On average, the common factor accounted for 22% of the total variance in the substance abuse scores. The substance-specific factors explained 34% of the variance. The average occasion-specific variance explained 15% of the variance (eTable 4, SI).

**Figure 2.**
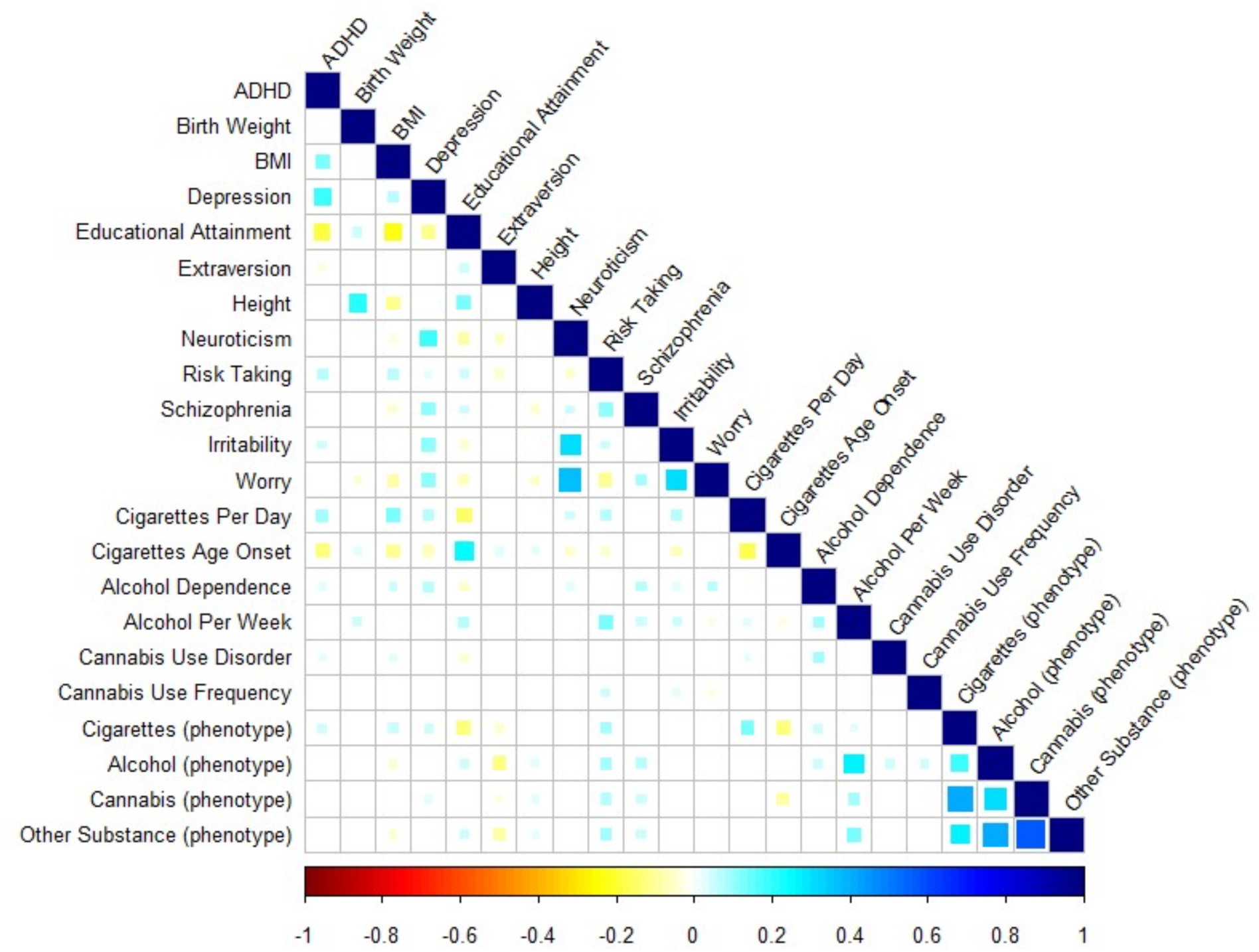
Correlations between the 18 PGSs and the mean scores of the substance abuse measures (cigarettes, alcohol, cannabis, and other substance) across age 17, 20 and 22. Data source: ALSPAC, N = 4218. ADHD = Attention deficit hyperactivity disorder. BMI = Body mass index. Blank cells represent non-significant coefficients (p > 0.05). The correlation estimates and p-values are reported in the SI (eTable).

### Effects of the PGSs reflecting substance abuse

The associations between the substance use PGSs with the common and substance-specific factors are shown in Table 2 and 3. As expected, the factors capturing cigarette and alcohol abuse were predicted by their respective PGSs (e.g. frequency of cigarette/alcohol use), reflecting specific genetic effects (e.g. linked to substance-specific metabolism). The common factor was independently predicted by two substance use PGSs (age of onset of cigarette use, alcohol frequency), in line with evidence implicating age of onset of cigarette use as a liability marker for initiation of use of other substances^40^. Other substance-specific factors were not predicted by their respective PGSs (e.g. cannabis abuse factor). This could reflect the fact that the GWAS used to derive those PGSs are only of limited power and have not yet succeeded in identifying genetic variants that are substance-specific in their biological function (e.g. metabolism)^41^.

### Effects of the PGSs reflecting vulnerabilities and protective traits

#### Common factor of substance abuse

In the single-PGS TSO models, three PGSs (risk taking, low extraversion, schizophrenia) were associated with the common factor of substance abuse after FDR correction and when included in the multi-PGSs TSO model (Table 2, Table 3, Figure 3). In the multi-PGSs model, the PGS for risk taking exerted the largest independent effect (b_standardized_=0.136, *p*_FDR_<0.001), followed by the PGS indexing low extraversion (b_standardized_=-0.095, *p*_FDR_<0.001) and schizophrenia (b_standardized_=0.056, *p*_FDR_=0.003).

**Figure 3.**
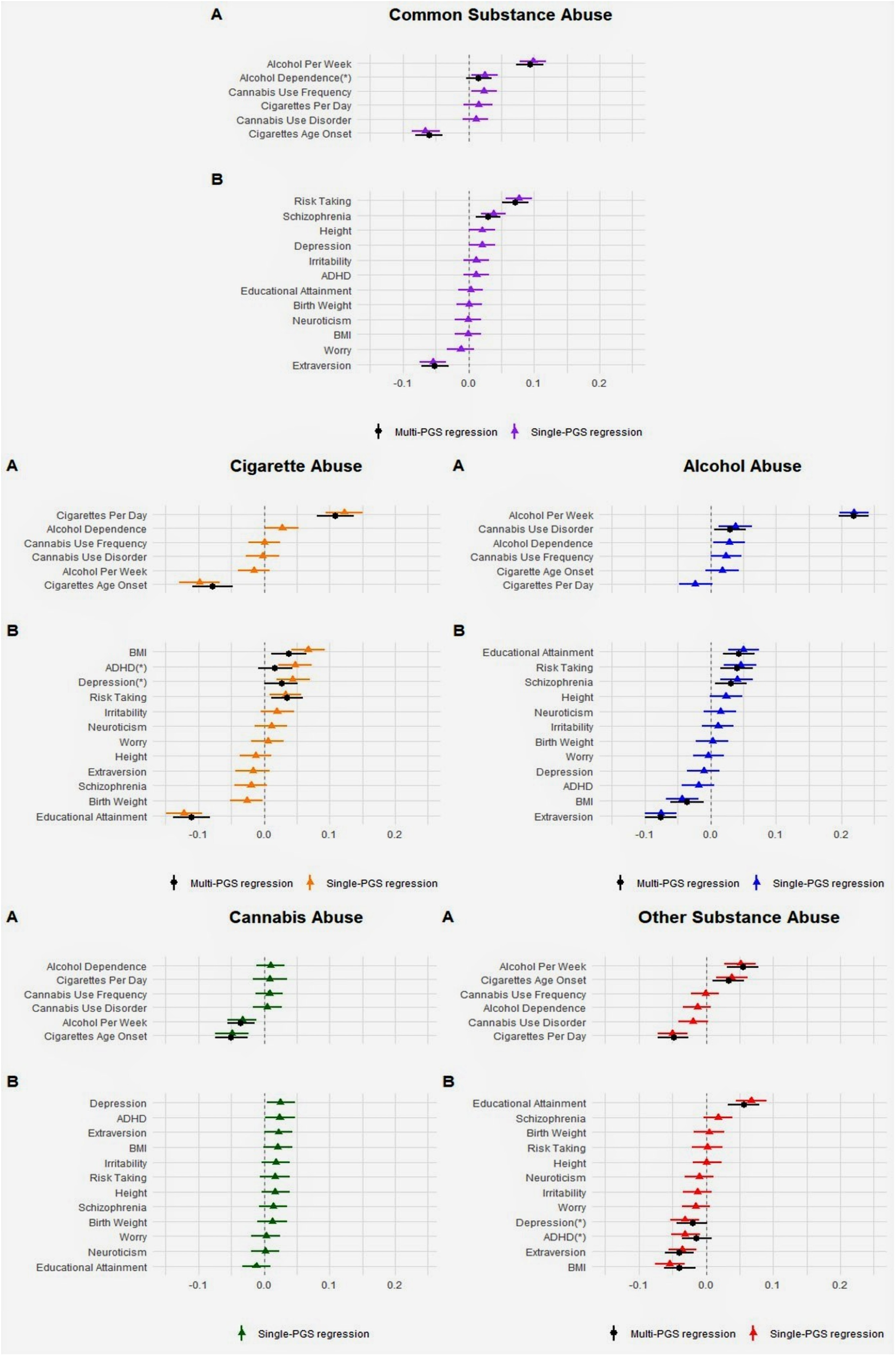
Single-PGS and multi-PGSs TSO models for the Common Substance, Cigarette, Alcohol, Cannabis and Other Substance Abuse factors. **Note.** Data source = ALSPAC, N = 4218. Model A = Substance Abuse PGSs. Model B = Individual Vulnerability and Protective Factors PGSs. TSO = Trait-State-Occasion. PGS = Polygenic Score. ADHD = Attention deficit hyperactivity disorder. BMI = Body Mass Index.

#### Substance-specific factor: Cigarette abuse

In the single-PGS TSO models, five PGSs were associated with the cigarette abuse factor following FDR correction (low educational attainment, BMI, ADHD, depression, risk taking). In the multi-PGSs TSO model, three PGSs remained associated with the cigarette abuse factor, including low educational attainment (b_standardized_= −0.151, *p*_FDR_<0.001) with the largest effect, followed by BMI (b_standardized_=0.052, *p*_FDR_=0.007) and risk taking (b_standardized_=0.048, *p*_FDR_=0.006).

#### Substance-specific factor: Alcohol abuse

In the single-PGS TSO models, five PGSs were associated with the alcohol abuse factor (low extraversion, educational attainment, risk taking, low BMI, schizophrenia), all of which remained significant following FDR correction and in the multi-PGSs TSO model. The largest effect was found for low extraversion (b_standardized_= −0.118, p *p*_FDR_<0.001), followed by educational attainment (b_standardized_=0.068, *p*_FDR_<0.001), risk taking (b_standardized_=0.063, *p*_FDR_=0.002), low BMI (b_standardized_= −0.055, *p*_FDR_=0.009) and schizophrenia (b_standardized_=0.049, *p*_FDR_=0.014).

#### Substance-specific factor: Cannabis abuse

No PGS was associated with the cannabis abuse factor.

#### Substance-specific factor: Other illicit substance abuse

In the single-PGS TSO models, five PGSs were associated with the factor representing other illicit substance abuse following FDR correction (educational attainment, low BMI, low extraversion, low depression, low ADHD). In the multi-PGSs TSO model, three PGSs remained independently associated, including educational attainment (b_standardized_=0.121, *p*_FDR_<0.001), low extraversion (b_standardized_= −0.085, *p*_FDR_<0.001) and low BMI (b_standardized_= −0.084, *p*_FDR=_0.002).

## DISCUSSION

This study is the first genomic investigation using the polygenic score (PGS) approach to examine the contribution of a range of individual traits and vulnerabilities to both common and specific liabilities to substance abuse. We highlight two findings: first, our results implicate a number of mental health vulnerabilities and personality traits in the common liability to substance abuse – namely, high risk taking, low extraversion, and schizophrenia. Second, we identified a distinct set of risk factors that independently contributed to substance-specific liabilities, such as educational attainment and BMI. In the following section, we will discuss: (1) insights for the aetiology of substance abuse, (2) findings regarding the common liability, (3) substance-specific findings, (4) implications for the prevention and treatment of substance abuse, and (5) limitations of our approach.

### Insights for the aetiology of substance abuse

Based on genomic evidence, our results helped to tease apart some of the genetic predispositions that indirectly contribute to common and substance-specific liabilities to substance abuse. In particular, different sets of genetically influenced mental health vulnerabilities and traits are likely to be involved in common versus substance-specific liabilities. Importantly, all associations found in this study need to be conceptualised as indirect effects of genetically influenced traits and vulnerabilities. To illustrate, our findings implicate that a genetic liability to risk taking would lead to greater risk taking behaviour, which, in turn, increases an individual’s propensity to engage in substance use irrespectively of the class of the substance. The use of PGSs as genetic proxies for individual vulnerabilities and traits is conceptually similar to the Mendelian Randomization (MR) framework^25^, where genetic instruments are used to investigate causal associations between an exposure and an outcome.

### Risk and protective factors involved in the common liability to substance abuse

Our results confirm previous findings of a common liability that partly underlies the abuse of different classes of addictive substances, such as cigarettes, alcohol, cannabis and other illicit substances^2^. Regarding its origins, our findings reveal that a genetic liability to risk taking, low extraversion and schizophrenia contribute to the common liability to substance abuse. This corroborates previous phenotypic evidence which reported associations between substance abuse and similar traits and vulnerabilities^11,13,14,42^. Intriguingly, a genetic predisposition for risk taking was most robustly associated with a common liability to substance abuse, but only to a lesser extent with substance-specific liabilities (cf. next paragraph). This indicates that individuals susceptible to risk taking are more likely to abuse an array of different substances, irrespectively of their class. In a similar vein, genetic predisposition to low extraversion was strongly associated with the common liability to substance abuse, whereas its associations with substance-specific liabilities were weaker. Thus, high extraversion may protect against the abuse of various substances. Furthermore, the common liability was influenced by genetic risk for schizophrenia. Taken together, these findings validate the idea that the abuse of various substances may reflect a self-medication strategy for those individuals more vulnerable to psychopathology and maladaptive personality traits^43^. Moreover, they suggest that shared genetic effects among different substances of abuse are substantially polygenic in nature, involving many genetic variants exerting indirect and small effects (e.g. polygenic influence via risk taking). This may explain why previous GWAS have largely been unsuccessful in identifying genetic variants common to different psychoactive substances. Future large GWAS may therefore benefit from modelling a common liability to substance abuse, similar to recent genome-wide attempts aiming to identify common genetic variation underlying psychiatric traits^44-46^.

In summary, our findings confirm that the common liability to substance abuse stems in part from a genetic contribution^47,48^. By using the PGS approach, we identified genetically influenced traits (i.e. risk taking, extraversion) and mental health vulnerabilities (i.e. schizophrenia), which independently contribute to a common liability to substance abuse.

### Risk and protective factors involved in substance-specific liabilities

Our results also showed that a substantial proportion of the phenotypic variation in substance abuse could not be explained by a common liability. Using the PGSs approach to identify risk and protective factors involved in the substance-specific liabilities revealed three patterns of associations. First (a), we identified a set of factors that were linked to both the common liability to substance abuse, as well as substance-specific liabilities. Second (b), several factors were linked to substance-specific liabilities but did not contribute to the common liability. Third (c), some traits previously implicated in substance abuse were not associated with any of the substance-specific liabilities.

Regarding (a), we found that all factors involved in the common liability including a genetic predisposition for risk taking, low extraversion and schizophrenia also contributed to the liability to alcohol abuse. Hence, the aetiologies of these two liabilities (i.e. alcohol vs. common) are partly based on overlapping risk factors. At the same time (b), our results showed that two individual traits – BMI and educational attainment – were not linked to the common liability but predicted substance-specific liabilities. Interesting results emerged regarding the direction of the identified associations. For example, we found that a predisposition for high educational attainment increased the risk of alcohol and illicit substance abuse but reduced the risk of cigarette abuse. This is consistent with the notion that education makes people less likely to smoke cigarettes due to an increased knowledge of its adverse health consequences^49^. At the same time, greater education may provide more opportunities to consume alcohol and access other substances, as indicated by previous observational evidence^50,51^. Opposite effects were also present for BMI. Here, a genetic predisposition for high BMI increased the risk of cigarette abuse, while reducing the risk of alcohol and other illicit substance abuse. The same pattern of associations has been reported in observational studies. For example, compared to normal weight adolescents, obese adolescents were at reduced risk of alcohol and illicit substance abuse, but had an elevated risk of cigarette abuse^52^. As nicotine is known to suppress appetite, this may suggest that adolescents with a greater predisposition to high BMI could smoke more in an attempt to control their appetite^53^. On the other hand, since alcohol has a high calorie content, individuals genetically predisposed to high BMI may consume less alcohol in order to control their weight^54^.

Finally (c), some of the previously implicated risk factors (e.g. neuroticism, ADHD)^10,11,13^ were not associated with the common or substance-specific liabilities in our sample. First, this could reflect a lack of power of the PGSs used in the analysis. However, we used powerful PGSs (e.g. neuroticism, derived from a GWAS with N>160000) that have been shown to predict rare outcomes in comparable samples^55^. Second, some PGSs were associated with substance abuse liabilities only in less controlled models (e.g. ADHD and depression predicting illicit substance abuse only in single-PGS but not multi-PGSs models). In addition to power issues, this may indicate that the effects of ADHD/depression were explained by potentially co-occurring traits that we included in our multivariable models.

### Insights for substance abuse prevention and treatment

Our findings offer insights for the prevention and treatment of substance abuse. First, we identified a set of individual vulnerabilities and traits, namely risk taking, low extraversion and schizophrenia, which contributed to the general liability to substance abuse. Hence, prevention and treatment programs aiming to reduce substance abuse in adolescents may benefit from focusing on those vulnerabilities and traits. For example, there is promising evidence from randomised controlled trials showing reductions in substance use following interventions targeting abilities related to risk taking (e.g. self-regulation) in adolescents^56^. Our results also highlight the importance to target those individuals at greatest risk of developing a problematic pattern of substance use based on pre-existing vulnerabilities such as schizophrenia. Hence, in adolescence with prodromal symptoms, particular emphasis may need to be placed on the prevention of substance abuse. Finally, it is important to better understand the mechanisms underlying some of the substance-specific associations found in this study (e.g. high BMI as a risk factor for cigarette abuse) in order to design more effective prevention and intervention strategies.

### Limitations

Despite the advantages of the PGS approach over studies that rely solely on phenotypic data^57^, an important limitation is that it is not possible to assure that all underlying assumptions of MR are fulfilled. However, sensitivity analyses as part of MR methods can help to assess potential violations (e.g. certain forms of pleiotropy). Such analyses will be possible once GWAS summary statistics for our outcomes of interest (i.e. common and specific liabilities to substance use) are available.

### Conclusion

Our findings reveal that distinct sets of genetically influenced vulnerabilities and protective factors are likely to be involved in the common versus substance-specific liabilities to substance abuse. In particular, genetic predisposition to high risk taking, low extraversion, and schizophrenia may increase the individual’s susceptibility to the abuse of any type of substance. Additionally, we identified genetic predisposition to educational attainment and BMI as independent risk factors for multiple specific substances, although in opposite directions. Prevention and treatment programs in adolescents may benefit from focusing on these vulnerabilities and protective factors.

## Additional contributions

We are extremely grateful to all the families who took part in this study, the midwives for their help in recruiting them, and the whole ALSPAC team, which includes interviewers, computer and laboratory technicians, clerical workers, research scientists, volunteers, managers, receptionists and nurses.

## REFERENCES

1. Rehm J, Mathers C, Popova S, Thavorncharoensap M, Teerawattananon Y, Patra J. Global burden of disease and injury and economic cost attributable to alcohol use and alcohol-use disorders. Lancet 2009; 373: 2223–2233.

2. Lynskey MT, Fergusson DM, Horwood LJ. The Origins of the Correlations between Tobacco, Alcohol, and Cannabis Use During Adolescence. J Child Psychol Psychiatry 1998; 39: 995–1005.

3. Agrawal A, Neale MC, Prescott CA, Kendler KS. Cannabis and other illicit drugs: Comorbid use and abuse/dependence in males and females. Behav Genet 2004; 34: 217–228.

4. John WS, Zhu H, Mannelli P, Schwartz RP, Subramaniam GA, Wu L-T. Prevalence, patterns, and correlates of multiple substance use disorders among adult primary care patients. Drug Alcohol Depend 2018; 187: 79–87.

5. DuPont RL, Han B, Shea CL, Madras BK. Drug use among youth: National survey data support a common liability of all drug use. Prev Med (Baltim) 2018; 187: 68–73.

6. Morley KI, Lynskey MT, Moran P, Borschmann R, Winstock AR. Polysubstance use, mental health and high-risk behaviours: Results from the 2012 Global Drug Survey. Drug Alcohol Rev 2015; 34: 427–437.

7. Bhalla IP, Stefanovics EA, Rosenheck RA. Clinical Epidemiology of Single Versus Multiple Substance Use Disorders. Med Care 2017; 55: S24–S32.

8. John WS, Zhu H, Mannelli P, Schwartz RP, Subramaniam GA, Wu L-T. Prevalence, patterns, and correlates of multiple substance use disorders among adult primary care patients. Drug Alcohol Depend 2018; 187: 79–87.

9. Morral AR, McCaffrey DF, Paddock SM. Reassessing the marijuana gateway effect. Addiction 2002; 97: 1493–1504.

10. Torrens M, Mestre-Pintó J-I, Domingo-Salvany A. Comorbidity of substance use and mental disorders in Europe. 2015. http://www.emcdda.europa.eu/system/files/publications/1988/TDXD15019ENN.pdf.

11. Swendsen J, Conway KP, Degenhardt L, Glantz M, Jin R, Merikangas KR et al. Mental disorders as risk factors for substance use, abuse and dependence: results from the 10-year follow-up of the National Comorbidity Survey. Addiction 2010; 105: 1117–1128.

12. Kwapil TR. A longitudinal study of drug and alcohol use by psychosis-prone and impulsive-nonconforming individuals. J Abnorm Psychol 1996; 105: 114–123.

13. Kotov R, Gamez W, Schmidt F, Watson D. Linking ‘Big’ personality traits to anxiety, depressive, and substance use disorders: A meta-analysis. Psychol Bull 2010; 136: 768–821.

14. Feldstein SW, Miller WR. Substance use and risk-taking among adolescents. J Ment Heal 2006; 15: 633–643.

15. Martínez-Loredo V, Fernández-Hermida JR, La Torre-Luque A de, Fernández-Artamendi S. Polydrug use trajectories and differences in impulsivity among adolescents. Int J Clin Heal Psychol 2018; 18: 235–244.

16. Erickson J, El-Gabalawy R, Palitsky D, Patten S, Mackenzie CS, Stein MB et al. Educational attainment as a protective factor for psychiatric disorders: Findings from a nationally representative longitudinal study. Depress Anxiety 2016; 33: 1013–1022.

17. Kanazawa S, Hellberg JEEU. Intelligence and Substance Use. Rev Gen Psychol 2010; 14: 382–396.

18. Kendler KS, Jacobson KC, Prescott CA, Neale MC. Specificity of genetic and environmental risk factors for use and abuse/dependence of cannabis, cocaine, hallucinogens, sedatives, stimulants, and opiates in male twins. Am J Psychiatry 2003; 160: 687–695.

19. Nivard MG, Verweij KJH, Minică CC, Treur JL, Vink JM, Boomsma DI. Connecting the dots, genome-wide association studies in substance use. Mol Psychiatry 2016; 21: 733–735.

20. Liu M, Jiang Y, Wedow R, Li Y, Brazel DM, Chen F et al. Association studies of up to 1.2 million individuals yield new insights into the genetic etiology of tobacco and alcohol use. Nat Genet 2019; 51: 237–244.

21. Furberg H, Kim Y, Dackor J, Boerwinkle E, Franceschini N, Ardissino D et al. Genome-wide meta-analyses identify multiple loci associated with smoking behavior. Nat Genet 2010; 42: 441–447.

22. Neale Lab. UK Biobank GWAS results. 2018.http://www.nealelab.is/uk-biobank.

23. Hancock DB, Reginsson GW, Gaddis NC, Chen X, Saccone NL, Lutz SM et al. Genome-wide meta-analysis reveals common splice site acceptor variant in CHRNA4 associated with nicotine dependence. Transl Psychiatry 2015; 5: e651–e651.

24. Euesden J, Lewis CM, O’Reilly PF. PRSice: Polygenic Risk Score software. Bioinformatics 2015; 31: 1466–1468.

25. Gage SH, Davey Smith G, Ware JJ, Flint J, Munafò MR. G = E: What GWAS Can Tell Us about the Environment. PLOS Genet 2016; 12: e1005765.

26. Pingault J-B, Murray J, Munafo M, Viding E. Causal inference in psychopathology: Using Mendelian randomisation to identify environmental risk factors for psychopathology. Psychopathol Rev 2016; a4: 4–25.

27. Verweij KJH, Abdellaoui A, Nivard MG, Sainz Cort A, Ligthart L, Draisma HHM et al. Short communication: Genetic association between schizophrenia and cannabis use. Drug Alcohol Depend 2017; 171: 117–121.

28. Hartz SM, Horton AC, Oehlert M, Carey CE, Agrawal A, Bogdan R et al. Association Between Substance Use Disorder and Polygenic Liability to Schizophrenia. Biol Psychiatry 2017; 82: 709–715.

29. Du Rietz E, Coleman J, Glanville K, Choi SW, O’Reilly PF, Kuntsi J. Association of Polygenic Risk for Attention-Deficit/Hyperactivity Disorder With Co-occurring Traits and Disorders. Biol Psychiatry Cogn Neurosci Neuroimaging 2018; 3: 635–643.

30. Carey CE, Agrawal A, Bucholz KK, Hartz SM, Lynskey MT, Nelson EC et al. Associations between polygenic risk for psychiatric disorders and substance involvement. Front Genet 2016; 7: 1–10.

31. Fraser A, Macdonald-Wallis C, Tilling K, Boyd A, Golding J, Davey Smith G et al. Cohort Profile: The Avon Longitudinal Study of Parents and Children: ALSPAC mothers cohort. Int J Epidemiol 2013; 42: 97–110.

32. Fagerström KO, Heatherton TF, Kozlowski LT. Nicotine addition and its assessment. Ear, nose, throat J 1990; 69: 763–765.

33. Saunders JB, Aasland OG, Babor TF, de la Fuente JR, Grant M. Development of the Alcohol Use Disorders Identification Test (AUDIT): WHO Collaborative Project on Early Detection of Persons with Harmful Alcohol Consumption. Addiction 1993; 88: 791–804.

34. Legleye S, Karila L, Beck F, Reynaud M. Validation of the CAST, a general population Cannabis Abuse Screening Test. J Subst Use 2007; 12: 233–242.

35. Rosseel Y. lavaan: An R Package for Structural Equation Modeling. J Stat Softw 2012; 48: 1–36.

36. Cole DA. Coping with longitudinal data in research on developmental psychopathology. Int J Behav Dev 2006; 30: 20–25.

37. Prenoveau JM. Specifying and Interpreting Latent State–Trait Models With Autoregression: An Illustration. Struct Equ Model 2016; 23: 731–749.

38. Newsom JT. Longitudinal Structural Equation Modeling. Taylor & Francis: New York, 2015.

39. Benjamini Y, Drai D, Elmer G, Kafkafi N, Golani I. Controlling the false discovery rate in behavior genetics research. Behav Brain Res 2001; 125: 279–84.

40. Agrawal A, Grant JD, Waldron M, Duncan AE, Scherrer JF, Lynskey MT et al. Risk for initiation of substance use as a function of age of onset of cigarette, alcohol and cannabis use: Findings in a Midwestern female twin cohort. Prev Med (Baltim) 2006; 43: 125–128.

41. Sherva R, Wang Q, Kranzler H, Zhao H, Koesterer R, Herman A et al. Genome-wide association study of cannabis dependence severity, novel risk variants, and shared genetic risks. JAMA Psychiatry 2016; 73: 472.

42. Volkow ND. Substance use disorders in schizophrenia--clinical implications of comorbidity. Schizophr Bull 2009; 35: 469–72.

43. Crum RM, Mojtabai R, Lazareck S, Bolton JM, Robinson J, Sareen J et al. A Prospective Assessment of Reports of Drinking to Self-medicate Mood Symptoms With the Incidence and Persistence of Alcohol Dependence. JAMA Psychiatry 2013; 70: 718.

44. Lee PH, Anttila V, Won H, Feng Y-CA, Rosenthal J, Zhu Z et al. Genome wide meta-analysis identifies genomic relationships, novel loci, and pleiotropic mechanisms across eight psychiatric disorders. bioRxiv 2019. doi:https://doi.org/10.1101/528117.

45. Grotzinger AD, Rhemtulla M, de Vlaming R, Ritchie SJ, Mallard TT, Hill WD et al. Genomic SEM provides insights into the multivariate genetic architecture of complex traits. bioRxiv 2018. doi:10.1101/305029.

46. Luningham JM, Poore HE, Yang J, Waldman ID. Testing structural models of psychopathology at the genomic level. bioRxiv 2018. doi:10.1101/502039.

47. Vanyukov MM, Tarter RE, Kirillova GP, Kirisci L, Reynolds MD, Kreek MJ et al. Common liability to addiction and “gateway hypothesis”: Theoretical, empirical and evolutionary perspective. Drug Alcohol Depend 2012; 123: S3–S17.

48. Kendler KS, Chen X, Dick D, Maes H, Gillespie N, Neale MC et al. Recent advances in the genetic epidemiology and molecular genetics of substance use disorders. Nat Neurosci 2012; 15: 181–189.

49. Office for National Statistics. Adult smoking habits in the UK: 2015. 2017 doi:10.2105/AJPH.76.11.1337.

50. Degenhardt L, Hall W. Extent of illicit drug use and dependence, and their contribution to the global burden of disease. Lancet 2012; 379: 55–70.

51. Huerta MC, Borgonovi F. Education, alcohol use and abuse among young adults in Britain. Soc Sci Med 2010; 71: 143–151.

52. Gearhardt AN, Waller R, Jester JM, Hyde LW, Zucker RA. Body mass index across adolescence and substance use problems in early adulthood. Psychol Addict Behav 2018; 32: 309–319.

53. Fulkerson J, French S. Cigarette smoking for weight loss or control among adolescents: gender and racial/ethnic differences. J Adolesc Heal 2003; 32: 306–313.

54. Traversy G, Chaput J-P. Alcohol Consumption and Obesity: An Update. Curr Obes Rep 2015; 4: 122–30.

55. Li JJ, Hilton EC, Lu Q, Hong J, Greenberg JS, Mailick MR. Validating psychosocial pathways of risk between neuroticism and late life depression using a polygenic score approach. J Abnorm Psychol 2019. doi:10.1037/abn0000419.

56. Pandey A, Hale D, Das S, Goddings A-L, Blakemore S-J, Viner RM. Effectiveness of Universal Self-regulation–Based Interventions in Children and Adolescents. JAMA Pediatr 2018; 172: 566.

57. Pingault J-B, O’Reilly PF, Schoeler T, Ploubidis GB, Rijsdijk F, Dudbridge F. Using genetic data to strengthen causal inference in observational research. Nat Rev Genet 2018; 19: 566–580.

