## Supplementary Material for "Identifying risk factors involved in the common versus specific liabilities to substance abuse: A genetically informed approach"

**Supplementary Information**

### eMETHODS

#### Substance Abuse Measures

*Cigarette Abuse*

Cigarette abuse was measured using the Fagerstrom Test[1], a validated measure of physical dependence on cigarette smoking which has been widely used across different countries in both clinical and non-clinical settings. The questionnaire consists of six items assessing the frequency and compulsion of cigarette consumption as well as nicotine dependence. The total Fagerstrom score can range between 0 and 10. For descriptive purposes (Table 1), we categorised the total score into three categories: low abuse (0-2), moderate abuse (3-5), and high abuse (6-10)[2].

*Alcohol Abuse*

Alcohol abuse was ascertained using the Alcohol Use Disorders Identification Test (AUDIT)[3]. This is a 10-item screening tool developed by the World Health Organisation to assess recent alcohol consumption, alcohol dependence symptoms, and alcohol-related problems. The AUDIT has been shown to provide a valid and reliable measure of alcohol abuse across gender, age, and cultures[4]. The total AUDIT score ranges from 0 to 40, with higher scores indicating greater alcohol abuse. In the descriptive analysis (Table 1), the total score was categorised into three groups: low abuse (0-8); moderate abuse (8-15); high abuse (16+)[4].

*Cannabis Abuse*

Cannabis abuse was assessed with the Cannabis Abuse Screening Test (CAST)[5], a validated tool for the screening of cannabis use disorders among adolescents and young adults. The questionnaire consists of six items pertaining to different aspects of problematic cannabis use such as non-recreational use (smoking alone or before midday), memory disorders, unsuccessful attempts to quit, and problems linked to cannabis consumption. The overall CAST score in ALSPAC ranges between 0 and 24. For descriptive purposes (Table 1), the total score was categorised into three groups: low abuse (1-8), moderate abuse (9-16), and high abuse (17-24)[5].

*Other illicit substances*

No validated questionnaire was available for the use of other illicit substances than cannabis. We therefore created an indicator representing the total number of other illicit substances used in the previous 12 months. These included: cocaine, amphetamines, inhalants, sedatives, hallucinogens, and opioids. The total score (1-6) was categorised into three groups: low abuse (1-2), moderate abuse (3-4), and high abuse (5-6).

**Analytical Strategy**

*Trait-State-Occasion (TSO) models.* The TSO models were tested using the specification model provided in Newsom[6]. The total scores of the four substance abuse measures at each time point were used as observed variables of three state factors, which in turn were indicators of a latent common substance abuse factor. The model included a separate occasion factor for the residual variance remaining after the trait factor variance was accounted for. In addition, four substance-specific (i.e. method) factors were specified accounting for the unique variance in each substance abuse measure across the three time points (eFigure 1). The adequacy of the TSO models was evaluated using Chi-Square Test ($\chi2$), Root Mean Square Error of Approximation (RMSEA), Comparative Fit Index (CFI), and Standardized Root-Mean-Square Residual (SRMR). $\chi^2$, RMSEA, and SRMR are all measures of absolute fit. Acceptable model fit is indicated by a statistically significant $\chi^2$ and RMSEA/ SRMR values less than .05. CFI is a comparative fit index which should be greater than .90[7].

**eFigure 1. Full TSO specification model.**

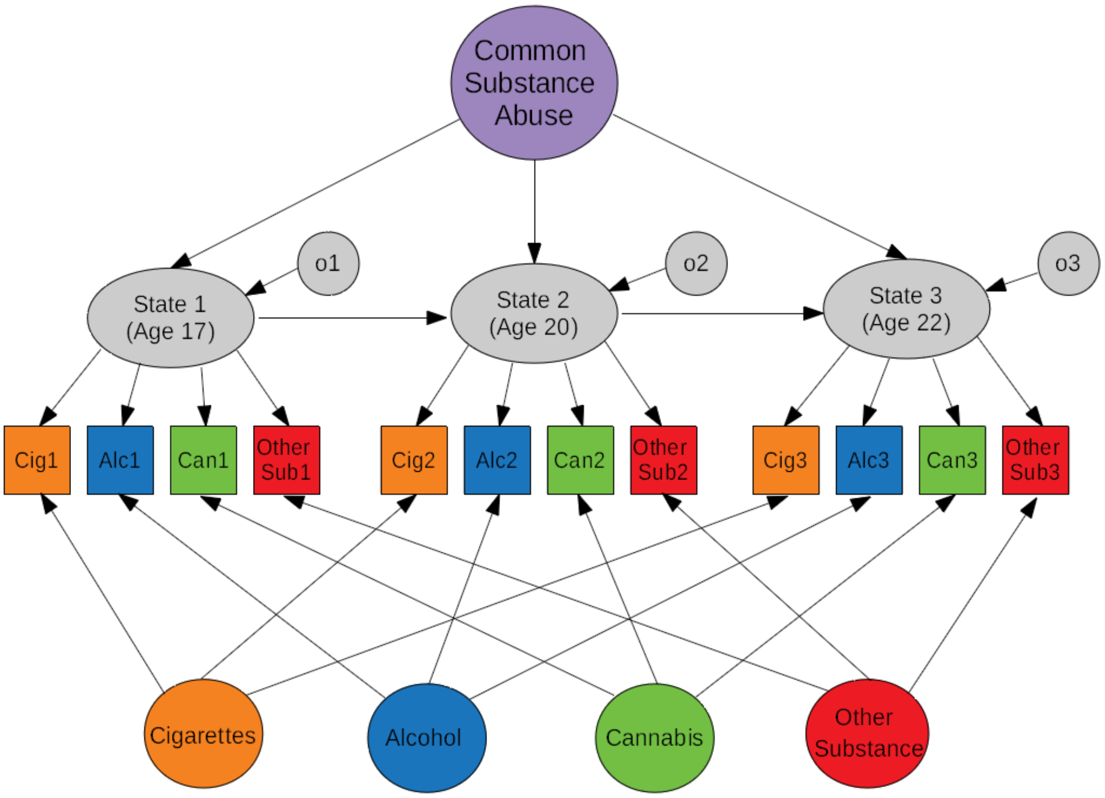


**Note**. The squares represent observed measures whereas the circles and elliptical shapes represent factors. The factors at the bottom represent substance-specific factors. o = occasion factor.

**eTable 1. Early sample characteristics for excluded versus included participants.**

| | Non-included | | | Included | | | Group comparisons | | |
|---|---|---|---|---|---|---|---|---|---|
| | *Mean* | *SD* | *N* | *Mean* | *SD* | *N* | *p-value*[*] | *r*[*] | *N* |
| Sex (% female)[a] | 43.9 | | 7690 | 56.9 | | 4218 | <.001 | 0.198 | 11908 |
| Parental divorce (%) yes)[a] | 17.7 | | 7647 | 11.5 | | 4192 | <.001 | 0.162 | 11839 |
| Parental mental illness (% | 4.0 | | 7647 | 4.8 | | 4192 | 0.046 | -0.052 | 11839 |
| Social class[b] | 3.29 | 1.23 | 6227 | 2.9 | 1.14 | 3728 | <.001 | -0.217 | 9955 |
| Birthweight (g) | 3420.98 | 545.63 | 7614 | 3443.55 | 513.55 | 4188 | 0.025 | 0.023 | 11802 |
| Mood score[c] | 15.97 | 6.03 | 5900 | 16.34 | 5.69 | 3748 | 0.002 | 0.039 | 9648 |
| Social achievement[d] score[d] | 19.16 | 3.93 | 6200 | 19.21 | 3.74 | 4046 | 0.517 | 0.005 | 10246 |

**Note**. [*]*p-value* estimates from significance tests, including t-tests (continuous variables) or chi-square tests (binary variables). *r* coefficients obtained from Spearman's Rho correlation tests. [a]Reported in % for binary variable. [b] Based on occupation, higher scores indicating lower social class. [c] Completed by mother (age: 6 months).[d] Completed by mother (age: 18 months). SD = standard deviation.

**eTable 2. Overview GWAS summary statistics.**

| Trait | Sample Size | Year published | Reference |
|---|---|---|---|
| ADHD | 55,374 | 2017 | Demontis D, Walters RK, Martin J, et al. Discovery of the first genome-wide significant risk loci for ADHD. BioRxiv. 2017:145581. |
| Agreeableness | 17,375 | 2012 | de Moor MHM, Costa PT, Terracciano A, et al. Meta-analysis of genome-wide association studies for personality. Mol Psychiatry. 2012;17(3):337-349. doi:10.1038/mp.2010.128. |
| Alcohol addiction ever | 6,514 | 2018 | Neale Lab. Uk Biobnak GWAS results - March 2018. http://www.nealelab.is/uk-biobank. Published |
| Alcohol Dependence | 38,686 | 2018 | Walters, R. K., Polimanti, R., Johnson, E. C., McClintick, J. N., Adams, M. J., Adkins, A. E., … Team, 23andMe Research. (2018). Transancestral GWAS of alcohol dependence reveals common genetic underpinnings with psychiatric disorders. Nature Neuroscience, 21(12), 1656–1669. https://doi.org/10.1038/s41593-018-0275-1 |
| Alcohol per week | 941,280 | 2019 | Liu, M., Jiang, Y., Wedow, R., Li, Y., Brazel, D. M., Chen, F., … Vrieze, S. (2019). Association studies of up to 1.2 million individuals yield new insights into the genetic etiology of tobacco and alcohol use. Nature Genetics. https://doi.org/10.1038/s41588-018-0307-5 |
| Anorexia nervosa | 14,477 | 2017 | Duncan L, Yilmaz Z, Gaspar H, et al. Significant locus and metabolic genetic correlations revealed in genome-wide association study of anorexia nervosa. Am J Psychiatry. 2017;174(9):850-858. doi:10.1176/appi.ajp.2017.16121402. |
| Anxiety | 18,186 | 2016 | Otowa T, Hek K, Lee M, et al. Meta-analysis of genome-wide association studies of anxiety disorders. Mol Psychiatry. 2016;21(10):1391-1399. doi:10.1038/mp.2015.197. |
| Autism spectrum disorder | 15,954 | 2017 | The Autism Spectrum Disorders Working Group of The Psychiatric Genomics Consortium. Meta-analysis of GWAS of over 16,000 individuals with autism spectrum disorder highlights a novel locus at 10q24.32 and a significant overlap with schizophrenia. Mol Autism . 2017;8(1):21. doi:10.1186/s13229-017-0137-9. |
| Bipolar disorder | 16,731 | 2011 | Psychiatric GWAS Consortium Bipolar Disorder Working Group. Large-scale genome-wide association analysis of bipolar disorder identifies a new susceptibility locus near ODZ4. Nat Genet. 2011;43(10):977-983. doi:10.1038/ng.943. |
| Birth weight | 205,475 | 2018 | Neale Lab. Uk Biobnak GWAS results - March 2018. http://www.nealelab.is/uk-biobank. Published 2018. |
| Body mass index | 681,275 | 2018 | Yengo L, Sidorenko J, Kemper KE, et al. Meta-analysis of genome-wide association studies for height and body mass index in~ 700,000 individuals of European ancestry. bioRxiv . 2018:274654. |
| Cannabis Use Disorder | 8,754 | 2016 | Sherva R, Wang Q, Kranzler H, et al. Genome-wide Association Study of Cannabis Dependence. JAMA Psychiatry.2016;73(5):472-480. doi: 10.1001/jamapsychiatry.2016.0036. Severity, Novel Risk Variants, and Shared Genetic Risks |
| Cannabis Use Frequency | 24,798 | 2018 | Neale Lab. Uk Biobnak GWAS results - March 2018. http://www.nealelab.is/uk-biobank. Published 2018. |
| Cigarette use (age of onset) | 341,427 | 2019 | Liu, M., Jiang, Y., Wedow, R., Li, Y., Brazel, D. M., Chen, F., … Vrieze, S. (2019). Association studies of up to 1.2 million individuals yield new insights into the genetic etiology of tobacco and alcohol use. Nature Genetics. https://doi.org/10.1038/s41588-018-0307-5 |
| Cigarette use (per day) | 337,334 | 2019 | Liu, M., Jiang, Y., Wedow, R., Li, Y., Brazel, D. M., Chen, F., … Vrieze, S. (2019). Association studies of up to 1.2 million individuals yield new insights into the genetic etiology of tobacco and alcohol use. Nature Genetics. https://doi.org/10.1038/s41588-018-0307-5 |
| Conscientiousness | 17,375 | 2012 | de Moor MHM, Costa PT, Terracciano A, et al. Meta-analysis of genome-wide association studies for personality. Mol Psychiatry . 2012;17(3):337-349. doi:10.1038/mp.2010.128. |
| Cross disorder | 61,220 | 2013 | Cross-Disorder Group of the Psychiatric Genomics Consotium. Identification of risk loci with shared effects on five major psychiatric disorders: a genome-wide analysis. Lancet . 2013;381(9875):1371-1379. doi:10.1016/S0140-6736(12)62129-1. |
| Depressive symptoms | 161,460 | 2016 | Okbay A, Baselmans BML, De Neve J-E, et al. Genetic variants associated with subjective well-being, depressive symptoms and neuroticism identified through genome-wide analyses. Nat Genet. 2016;48(6):624-633. doi:10.1038/ng.3552. |
| Diagnosis of depression | 173,005 | 2018 | Wray NR, Ripke S, Mattheisen M, et al. Genome-wide association analyses identify 44 risk variants and refine the genetic architecture of major depression. Nat Genet. 2018;50(5):668-681. doi:10.1038/s41588-018-0090-3. |
| Educational attainment | 766,345 | 2018 | Lee JJ, Wedow R, Okbay A, et al. Gene discovery and polygenic prediction from a genome-wide association study of educational attainment in 1.1 million individuals. Nat Genet. 2018;50(8):1112-1121. doi:10.1038/s41588-018-0147-3. |
| Extraversion | 17,375 | 2012 | de Moor MHM, Costa PT, Terracciano A, et al. Meta-analysis of genome-wide association studies for personality. Mol Psychiatry . 2012;17(3):337-349. doi:10.1038/mp.2010.128. |
| Extraversion (Item Response Theory) | 63,030 | 2015 | van den Berg SM, de Moor MHM, McGue M, et al. Harmonization of Neuroticism and Extraversion phenotypes across inventories and cohorts in the Genetics of Personality Consortium: an application of Item Response Theory. Behav Genet . 2014;44(4):295-313. doi:10.1007/s10519-014-9654-x. |
| Height | 693,529 | 2018 | Yengo L, Sidorenko J, Kemper KE, et al. Meta-analysis of genome-wide association studies for height and body mass index in~ 700,000 individuals of European ancestry. bioRxiv . 2018:274654. |
| Insomnia | 113,006 | 2017 | Hammerschlag AR, Stringer S, de Leeuw CA, et al. Genome-wide association analysis of insomnia complaints identifies risk genes and genetic overlap with psychiatric and metabolic traits. Nat Genet. 2017;49(11):1584-1592. doi:10.1038/ng.3888. |
| Intelligence | 269,867 | 2018 | Savage JE, Jansen PR, Stringer S, et al. Genome-wide association meta-analysis in 269,867 individuals identifies new genetic and functional links to intelligence. Nat Genet. 2018;50(7):912-919. doi:10.1038/s41588-018-0152-6. |

| | | | |
|---|---|---|---|
| Internalizing symptoms | 4,596 | 2014 | Benke KS, Nivard MG, Velders FP, et al. A genome-wide association meta-analysis of preschool internalizing problems. J Am Acad Child Adolesc Psychiatry. 2014;53(6):667-676.e7. doi:10.1016/j.jaac.2013.12.028. |
| Irritability | 345,231 | 2018 | Neale Lab. Uk Biobnak GWAS results - March 2018. http://www.nealelab.is/uk-biobank. Published 2018. |
| Loneliness | 10,760 | 2017 | Gao J, Davis LK, Hart AB, et al. Genome-wide association study of loneliness demonstrates a role for common variation. Neuropsychopharmacology . 2017;42(4):811-821. doi:10.1038/npp.2016.197. |
| Neuroticism | 168,105 | 2018 | Turley P, Walters RK, Maghzian O, et al. Multi-trait analysis of genome-wide association summary statistics using MTAG. Nat Genet. 2018;50(2):229-237. doi:10.1038/s41588-017-0009-4. |
| Obsessive compulsive disorder | 9,725 | 2018 | Arnold PD, Askland KD, Barlassina C, et al. Revealing the complex genetic architecture of obsessive–compulsive disorder using meta-analysis. Mol Psychiatry . 2018;23(5):1181-1188. doi:10.1038/mp.2017.154. |
| Openness | 17,375 | 2012 | de Moor MHM, Costa PT, Terracciano A, et al. Meta-analysis of genome-wide association studies for personality. Mol Psychiatry . 2012;17(3):337-349. doi:10.1038/mp.2010.128. |
| Risk taking | 436,236 | 2018 | Clifton EAD, Perry JRB, Imamura F, et al. Genome–wide association study for risk taking propensity indicates shared pathways with body mass index. Commun Biol. 2018;1(1):36. doi:10.1038/s42003-018-0042-6. |
| Schizophrenia | 105,318 | 2018 | Pardiñas AF, Holmans P, Pocklington AJ, et al. Common schizophrenia alleles are enriched in mutation-intolerant genes and in regions under strong background selection. Nat Genet. 2018;50(3):381-389. doi:10.1038/s41588-018-0059-2. |
| Worry | 348,219 | 2018 | Nagel M, Jansen PR, Stringer S, et al. Meta-analysis of genome-wide association studies for neuroticism in 449,484 individuals identifies novel genetic loci and pathways. Nat Genet. 2018;50(7):920-927. doi:10.1038/s41588-018-0151-7. |

**eTable 3. Summary of GWAS summary statistics excluded and included in the analysis.**

| **Excluded GWASs (Risk Factors)**<br><br>Exclusion criteria:<br>1 = Content Overlap;<br>2 = GWAS < 20,000 | **Reasons for exclusion** | **Included GWASs** |
|---|---|---|
| Cross Disorder | 1 | ***Risk Factors*** |
| Internalising | 1, 2 | ADHD |
| Loneliness | 2 | Birth weight |
| OCD | 2 | Body mass index |
| Anorexia | 2 | Depression |
| Autism | 2 | Educational attainment |
| Depression Symptoms | 1 | Extraversion (Item Response Theory) |
| Extraversion | 1, 2 | Height |
| Intelligence | 1 | Neuroticism |
| Agreeableness | 2 | Risk taking |
| Anxiety | 2 | Schizophrenia |
| Bipolar Disorder | 2 | Irritability |
| Conscientiousness | 2 | Worry |
| Openness | 2 | |
| Insomnia | 1 | ***Substance Abuse*** |
| | | Alcohol Dependence |
| | | Alcohol Per Week |
| | | Cigarettes Age Onset |
| | | Cigarettes Per Day |
| | | Cannabis Use Disorder |
| | | Cannabis Use Frequency |

**eTable 4. TSO model parameters.**

| | R-squared | R-squared decomposition | | |
|---|---|---|---|---|
| | | *SA Trait* | *Occasion-specific* | *Substance -specific* |
| *Age 17* | | | | |
| Cigarettes | 0.576 | 0.132 | 0.085 | 0.357 |
| Alcohol | 0.396 | 0.115 | 0.074 | 0.206 |
| Cannabis | 0.741 | 0.378 | 0.243 | 0.119 |
| Other substances | 0.555 | 0.308 | 0.198 | 0.047 |
| *Age 20* | | | | |
| Cigarettes | 0.954 | 0.115 | 0.094 | 0.744 |
| Alcohol | 0.789 | 0.104 | 0.085 | 0.599 |
| Cannabis | 0.869 | 0.336 | 0.275 | 0.257 |
| Other substances | 0.870 | 0.286 | 0.234 | 0.349 |
| *Age 22* | | | | |
| Cigarettes | 0.645 | 0.113 | 0.074 | 0.456 |
| Alcohol | 0.645 | 0.114 | 0.075 | 0.455 |
| Cannabis | 0.876 | 0.316 | 0.209 | 0.350 |
| Other substances | 0.677 | 0.293 | 0.194 | 0.190 |
| *Average R-squared* [illegible] | 72% | 22% | 15% | 34% |

**Note.** TSO model without risk factors. TSO = Trait-state-occasion. SA = substance abuse.

**eTable 5. Estimates of the correlation between the *18* PGSs and the mean scores of the measures of substance abuse (cigarettes, alcohol, cannabis, and other substances) across age 17, 20 and 22.**

| | *ADHD* | *Birth Weight* | *BMI* | *Depression* | *Educational Attainment* | *Extraversion* | *Height* | *Neuroticism* | *Risk Taking* | *Schizophrenia* | *Irritability* | *Worry* | *Cigarettes Per Day* | *Cigarettes Age Onset* | *Alcohol Dependence* | *Alcohol Per Week* | *Cannabis Use Disorder* | *Cannabis Use Frequency* | *Cigarettes (phenotype)* | *Alcohol (phenotype)* | *Cannabis (phenotype)* |
|---|---|---|---|---|---|---|---|---|---|---|---|---|---|---|---|---|---|---|---|---|---|
| ADHD | | | | | | | | | | | | | | | | | | | | | |
| Birth Weight | -0.01 | | | | | | | | | | | | | | | | | | | | |
| BMI | 0.13*** | 0.01 | | | | | | | | | | | | | | | | | | | |
| Depression | 0.20*** | -0.01 | 0.07*** | | | | | | | | | | | | | | | | | | |
| Educational Attainment | -0.19*** | 0.06*** | -0.22*** | -0.11*** | | | | | | | | | | | | | | | | | |
| Extraversion | -0.04* | 0.02 | -0.01 | -0.03 | 0.06*** | | | | | | | | | | | | | | | | |
| Height | -0.03 | 0.22*** | -0.11*** | -0.01 | 0.13*** | -0.02 | | | | | | | | | | | | | | | |
| Neuroticism | 0.03 | -0.03 | -0.04* | 0.18*** | -0.08*** | -0.07*** | -0.02 | | | | | | | | | | | | | | |
| Risk Taking | 0.07*** | 0.01 | 0.08*** | 0.04* | 0.05** | -0.05** | 0.00 | -0.05** | | | | | | | | | | | | | |
| Schizophrenia | 0.02 | 0.00 | -0.05** | 0.11*** | 0.04* | -0.02 | -0.05** | 0.04* | 0.11*** | | | | | | | | | | | | |
| Irritability | 0.04* | 0.01 | 0.02 | 0.11*** | -0.05*** | -0.02 | -0.02 | 0.31*** | 0.05** | 0.03 | | | | | | | | | | | |
| Worry | 0.00 | -0.04** | -0.10*** | 0.12*** | -0.07*** | 0.02 | -0.05** | 0.37*** | -0.11*** | 0.09*** | 0.30*** | | | | | | | | | | |
| Cigarettes Per Day | 0.10*** | -0.01 | 0.13*** | 0.07*** | -0.15*** | 0.00 | -0.02 | 0.04** | 0.07*** | 0.01 | 0.06*** | 0.01 | | | | | | | | | |
| Cigarettes Age Onset | -0.13*** | 0.04* | -0.10*** | -0.08*** | 0.25*** | 0.04* | 0.03* | -0.04** | -0.05** | 0.00 | -0.07*** | 0.00 | -0.18*** | | | | | | | | |
| Alcohol Dependence | 0.04* | 0.00 | 0.04** | 0.07*** | -0.04** | 0.02 | -0.02 | 0.04* | 0.03 | 0.07*** | 0.04* | 0.06*** | 0.01 | -0.03 | | | | | | | |
| Alcohol Per Week | -0.01 | 0.05** | -0.03 | 0.02 | 0.07*** | -0.03 | 0.01 | 0.01 | 0.12*** | 0.06*** | 0.05** | -0.03* | 0.03* | -0.04* | 0.08*** | | | | | | |
| Cannabis Use Disorder | 0.03* | 0.03 | 0.03* | 0.03 | -0.04** | -0.02 | 0.02 | 0.00 | 0.01 | -0.01 | 0.01 | -0.01 | 0.03* | -0.02 | 0.08*** | 0.03 | | | | | |
| Cannabis Use Frequency | 0.02 | -0.01 | -0.01 | 0.00 | 0.01 | 0.00 | 0.03 | -0.01 | 0.05** | 0.02 | 0.03* | -0.03* | 0.00 | -0.01 | -0.01 | 0.02 | 0.02 | | | | |
| Cigarettes (phenotype) | 0.05** | -0.02 | 0.06*** | 0.05** | -0.12*** | -0.05** | 0.00 | 0.01 | 0.08*** | 0.01 | 0.03 | 0.00 | 0.13*** | -0.13*** | 0.05** | 0.04* | 0.02 | 0.02 | | | |
| Alcohol (phenotype) | -0.01 | 0.00 | -0.05** | 0.01 | 0.05*** | -0.12*** | 0.04* | 0.02 | 0.08*** | 0.07*** | 0.02 | -0.01 | -0.02 | -0.02 | 0.05** | 0.26*** | 0.05** | 0.04** | 0.20*** | | |
| Cannabis (phenotype) | 0.02 | 0.01 | 0.02 | 0.04* | 0.00 | -0.03* | 0.03* | 0.00 | 0.08*** | 0.05*** | 0.02 | -0.01 | 0.02 | -0.09*** | 0.03 | 0.09*** | 0.01 | 0.03 | 0.42*** | 0.30*** | |
| Other Substance (phenotype) | -0.02 | 0.00 | -0.05** | 0.01 | 0.06*** | -0.08*** | 0.03* | -0.01 | 0.08*** | 0.06*** | 0.00 | -0.02 | -0.02 | -0.03 | 0.02 | 0.13*** | 0.00 | 0.02 | 0.26*** | 0.41*** | 0.57*** |

**Note**. p< .001 ***; p<.01 **; p<.05*.

**References**

1 Fagerström KO, Heatherton TF, Kozlowski LT. Nicotine addition and its assessment. *Ear, nose, throat J* 1990; **69**: 763–765.

2 National Institute of Drug Abuse. Instrument: Fagerstrom Test For Nicotine Dependence (FTND). https://cde.drugabuse.gov/instrument/d7c0b0f5-b865-e4de-e040-bb89ad43202b.

3 Saunders JB, Aasland OG, Babor TF, de la Fuente JR, Grant M. Development of the Alcohol Use Disorders Identification Test (AUDIT): WHO Collaborative Project on Early Detection of Persons with Harmful Alcohol Consumption. *Addiction* 1993; **88**: 791–804.

4 Lawford BR, Barnes M, Connor JP, Heslop K, Nyst P, Young RM. Alcohol Use Disorders Identification Test (AUDIT) scores are elevated in antipsychotic-induced hyperprolactinaemia. *J Psychopharmacol* 2012; **26**: 324–329.

5 Legleye S, Karila L, Beck F, Reynaud M. Validation of the CAST, a general population Cannabis Abuse Screening Test. *J Subst Use* 2007; **12**: 233–242.

6 Newsom JT. *Longitudinal Structural Equation Modeling*. Taylor & Francis: New York, 2015.

7 Byrne BM. *Structural equation modeling with Mplus: Basic concepts, applications, and programming*. Routledge/Taylor & Francis Group: New York, US, 2012.
